# Coincident bursts of high frequency oscillations across the human cortex coordinate large-scale memory processing

**DOI:** 10.1101/2025.03.24.644989

**Authors:** S. Prathapagiri, J. Cimbalnik, J.S. Garcia Salinas, M. Galanina, L. Jurkovicova, P. Daniel, M. Kojan, R. Roman, M. Pail, W. Fortuna, M. Sluzewska-Niedzwiedz, P. Tabakow, A. Czyzewski, M. Brazdil, M.T. Kucewicz

## Abstract

Oscillations in the high gamma and ripple frequency ranges are known to coordinate local hippocampal and neocortical neuronal assemblies during memory encoding and recall. Here, we explored spatiotemporal dynamics and the role of global coordination of these fast oscillatory discharges across the sensory and associational cortical areas in distinct phases of memory processing. Individual bursts of high frequency oscillations were detected in intracranial recordings from epilepsy patients remembering word lists for immediate free recall. We found constant coincident bursting across visual and higher order processing areas, peaking before recall and elevated during encoding of words. This global co-bursting was modulated by memory processing, engaged approximately half of the recorded electrode sites, and clustered into a sequence of multiple consecutive bursting discharges. Our results suggest a general role of global coincident high frequency oscillations in organizing large-scale information processing across the brain necessary especially, but not exclusively, for memory functions.

## Introduction

How widespread brain regions dynamically communicate to generate coherent behavior in response to external or internal stimuli remains difficult to capture with a high temporal resolution but limited spatial sampling of the current electrophysiological methods. Temporal coordination of neural activities has been proposed as a universal mechanism for integrating information across the brain ^1–3^. Oscillations in gamma frequency ranges beyond the classic Berger bands of the EEG spectrum (30 - 80Hz)^4^, were originally shown to synchronize neuronal firing responses to particular attended stimuli in the cat visual cortex ^5,6^, suggesting a role for high-frequency oscillatory activities in binding coherent object representations ^7^. Although the exact role in this perceptual binding is debatable ^8^, oscillations in high gamma and ripple frequency ranges (>60 Hz) have recently been revisited in the context of more general retrieving and integrating information in the brain. Ripple frequency oscillations across the hippocampus and neocortex are associated with encoding and recalling remembered stimuli ^9–15^. Several groups have now confirmed that these oscillations are coupled in time within less than 500 ms across particular functionally-related and widespread cortical areas ^11,12,16–18^, leading back to the hypothesis that such temporally coordinated high-frequency activities can facilitate integration and binding of information processing across the brain ^19^. This would provide an elegant mechanism for global communication through temporal coordination of these fast activities in distributed cortical areas without necessarily requiring precise phase synchronization.

High frequency oscillations (HFOs) present a plausible activity for large-scale tracking of coordinated electrophysiological discharges both locally at the level of neuronal assemblies and globally across networks of connected areas ^20^. HFOs comprise a wide range of physiological and epileptiform discharges spanning gamma, ripple and fast ripple frequency ranges ^20–27^. Originally reported as sharp-wave ripple complexes in the rodent hippocampus and associated with physiological functions ^28^, HFOs were subsequently found in cortical networks supporting memory and cognitive functions ^11–13,17,29,30^. In humans, they were first described as bursts of high gamma power in visual cortex ^31^, as ripple oscillations in the mesial temporal cortex during memory task performance ^32^, and across the cortex in the entire range of gamma, ripple and fast ripple frequencies (60-500 Hz) induced by memory encoding and recall ^13^. Although multiple terms and definitions have been established for ripple oscillations and other high frequency activities ^20–22,24^, there is a general consensus that they reflect coordinated firing of local neuronal ensembles that can be detected and tracked in time and anatomical space.

Growing evidence from multiple groups shows that physiological ripple frequency oscillations in the human cortex are underpinned by coordinated phase-locked spiking of neuronal assemblies ^9,11,17,33,34^. This neuronal spiking is not only locked to phases of the oscillation but also reveals specific firing sequences that can be used to decode particular words and predict their correct recall ^9^. The sequences are reactivated following presentation of word stimuli for encoding until their recall. This bursting of spike sequences across multiple neurons occurs at the times of HFO discharges, which in turn are detected around critical events like encoding or recalling stimuli ^10,12,13,16^. Multiple coincident HFOs (co-HFOs) have lately been shown to co-occur at the critical points of scene cuts during movie watching when a memory trace for a given scene that was just displayed is encoded and then replayed during subsequent co-HFO events ^15^. Altogether, these studies support a role of coordinated bursting of high frequency oscillations in integrating information for particular memory traces and thus can be used as neural substrates of engram activities ^20^, reflecting storage and recall of declarative memories.

In this study, we investigate this coincident HFO discharges in response to four types of events in a Free Recall verbal memory task related to preparation, encoding, distraction and recalling of stimuli to test its possible behavioral functions. Our hypothesis is that the global HFO bursting is modulated primarily by memory processing. We explore whether this global cortical coordination ^11^ engages mainly sensory or higher order processing areas of the association cortex, as suggested in the previous research ^12,15,16,18^. How selective are these globally coordinated oscillations in terms of specific stimuli remembered and recording sites involved? Animal research suggests that encoding a single engram can engage half of the brain areas studied ^35^. Finally, what is the temporal organization of distinct coincident HFO discharges? HFO bursting is known to increase before the onset of a behavioral response ^9,10,12,15,16^, but it is unclear how this is related to the actual moment of explicit or implicit recall. Whether this is underpinned by a single bursting event or a sequence of distinct cortical networks in a train of bursting events has been difficult to assess in the previous paradigms of freely recalling multiple stimuli and complex pictures or movie scenes together but possible with individual word concepts in this study. Our goal was to determine the large-scale structure of coincident HFO discharges underlying memory processing of common word concepts.

## Results

A total of 5,266,937 distinct bursts of HFOs with discrete duration of at least four cycles and peak frequency within a broad 50-600 Hz range ^13,14^ were detected from cortical stereo EEG depth electrode contacts implanted in epilepsy patients (Table 1) during a Free Recall (FR) task performance, and additional verbal memory tasks such as Paired-Associate Learning (PAL) ^36^. The HFO burst detections were made around four phases of the FR task: (1) countdown digits before the start of each trial, (2) presentation of words for memory encoding, (3) presentation of arithmetic problems during the delay period, and (4) onset of verbalization of the successfully recalled words (Fig. 1a), which was semi-automatically marked based on audio signal processing (Suppl. fig. S2). We confirmed previously reported findings of an increased rate of HFO bursts induced by memory encoding and especially by memory recall ^10,12,13^ (Fig. 1a,b). We observed this increased rate of HFO detections in the sensory visual areas of the occipital cortex both during visual presentation of word stimuli and at the time of free verbal recall of the remembered words in the absence of any visual stimulation on the computer screen (Fig. 1b,c). This internally-driven sensory activation in the occipital cortex may reflect mental imagery of the recalled words - patients were encouraged to visualize the remembered words to help them encode these more effectively.

**Figure 1.**
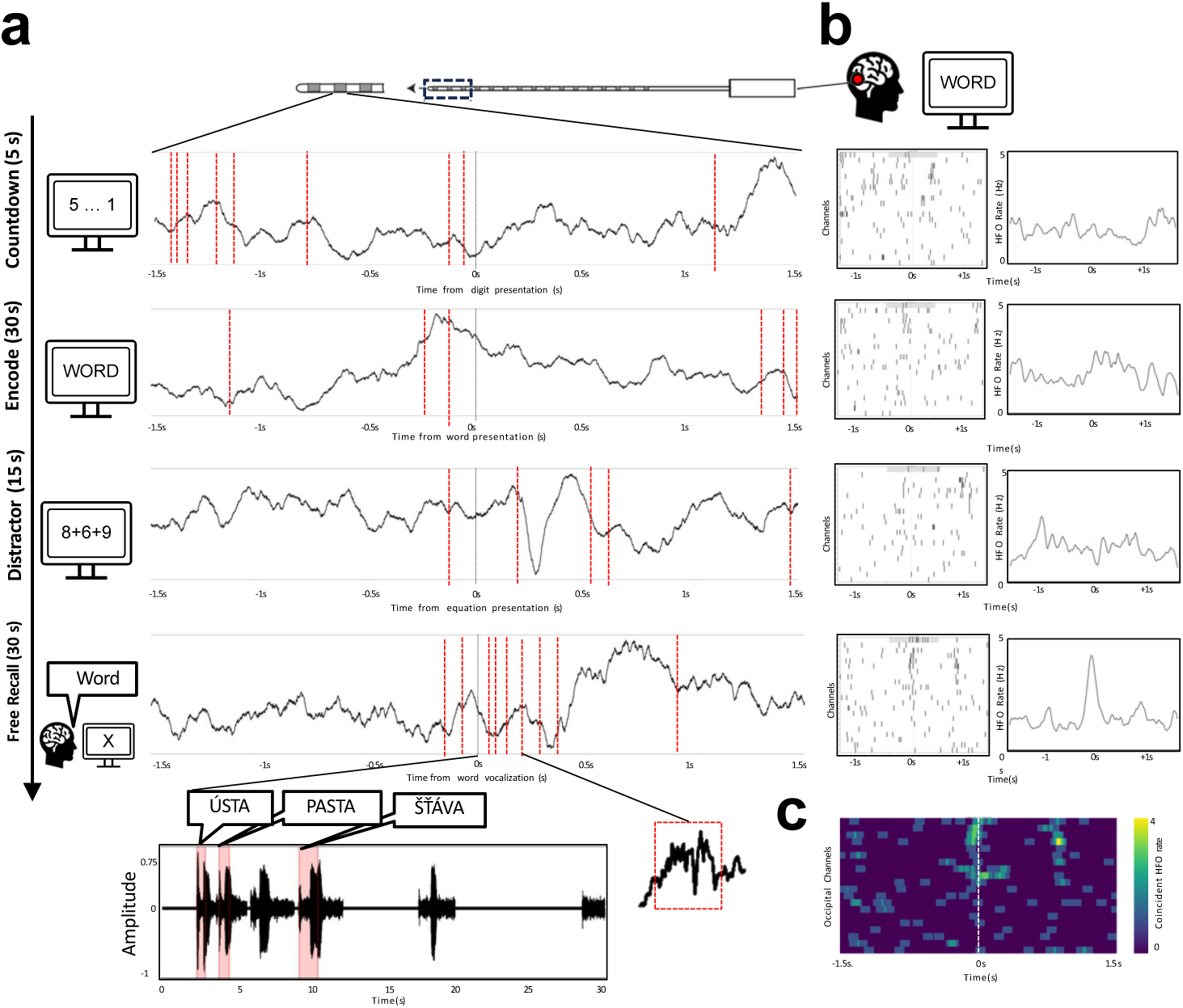
Bursts of high frequency oscillations co-occur across multiple cortical electrode contacts especially before recalling words. a) Raw LFP traces from an example macro-electrode contact implanted in the occipital cortex show timepoints when individual HFO bursts were detected (red lines) around the times of presenting a countdown digit, an encoded word, a distractor equation, and of recalling a word. b) HFOs detected across all channels implanted in the occipital cortex for example traces in ‘a’ with a summary of the mean detection rate from the entire session. c) Matrix of coincident detections in 100ms time bins from the example recall phase raster in ‘b’. Notice highly coincident bursting among a subgroup of these visual cortical electrodes even though no visual stimulation is delivered in this phase of the task - patients are freely recalling remembered words with the gaze fixated in the middle of a blank screen.

**Table 1.** Summary of Subject Demographics and Electrode Contact Data. Demographic and recording details of subjects included in the study, categorized by language (ISO 639-1 code; CS: Czech, SK: Slovak, PL: Polish).

| Dataset | Language | Subject ID | Sex | Age | FR runs | PAL runs | Micro contacts | Macro contacts |
| --- | --- | --- | --- | --- | --- | --- | --- | --- |
| BR | CS | 2 | F | 31 | 1 | 1 | 0 | 173 |
| BR | CS | 3 | M | 40 | 1 | 1 | 9 | 118 |
| BR | CS | 5 | M | 29 | 1 | 1 | 0 | 174 |
| BR | SK | 6 | M | 21 | 1 | 1 | 0 | 173 |
| BR | CS | 7 | M | 36 | 2 | 0 | 0 | 170 |
| BR | SK | 8 | M | 31 | 1 | 1 | 0 | 157 |
| BR | SK | 9 | M | 30 | 1 | 0 | 0 | 174 |
| BR | CS | 10 | F | 27 | 2 | 0 | 0 | 172 |
| BR | SK | 11 | F | 36 | 2 | 2 | 0 | 172 |
| BR | CS | 13 | M | 43 | 1 | 1 | 0 | 102 |
| BR | CS | 14 | M | 41 | 1 | 0 | 10 | 134 |
| BR | CS | 16 | F | 41 | 2 | 0 | 0 | 146 |
| WR | PL | 1 | F | 24 | 2 | 2 | 26 | 56 |
| WR | PL | 2 | M | 31 | 3 | 3 | 44 | 44 |
| WR | PL | 3 | F | 32 | 2 | 2 | 48 | 40 |
| WR | PL | 4 | M | 28 | 2 | 3 | 57 | 58 |
| WR | PL | 5 | F | 22 | 3 | 3 | 58 | 44 |
| Total counts | CS - 8, SK - 4, PL - 5 | 17 | 10 M | 31.9 | 34 | 24 | 282 | 2957 |

We quantified each HFO detection as a single point in time ^13,37^ taking the midpoint between burst onset and offset (Suppl. fig. S1) to study not only the rates within particular channels but also temporal correlations between channels from the same and different cortical regions. Despite the elevated rates recorded on selected electrode contacts at the encoding or recall phases of the task, there were no significant differences in overall background bursting rates across the four phases of the task (Kruskal-Wallis test, H(3) = 2.94, p = 0.40, N=12 ; Suppl. fig. S3). The baseline rates in particular frequency ranges were consistent with those reported in previous studies ^10,13,14^, which is approximately 0.8 bursts detected per second in the high gamma/ripple frequency range (60-250 Hz) per channel. Based on the findings of coupled hippocampal-cortical ^12^ and cortical-cortical ^11^ HFO detections, we sorted the channels by their pairwise correlation (Pearson’s) and found a pattern of coincident HFO (co-HFO) detections within 100 ms window across multiple channels (Fig. 1c), which was most prominent at the moment of recall onset (Fig. 2b-c).

**Figure 2.**
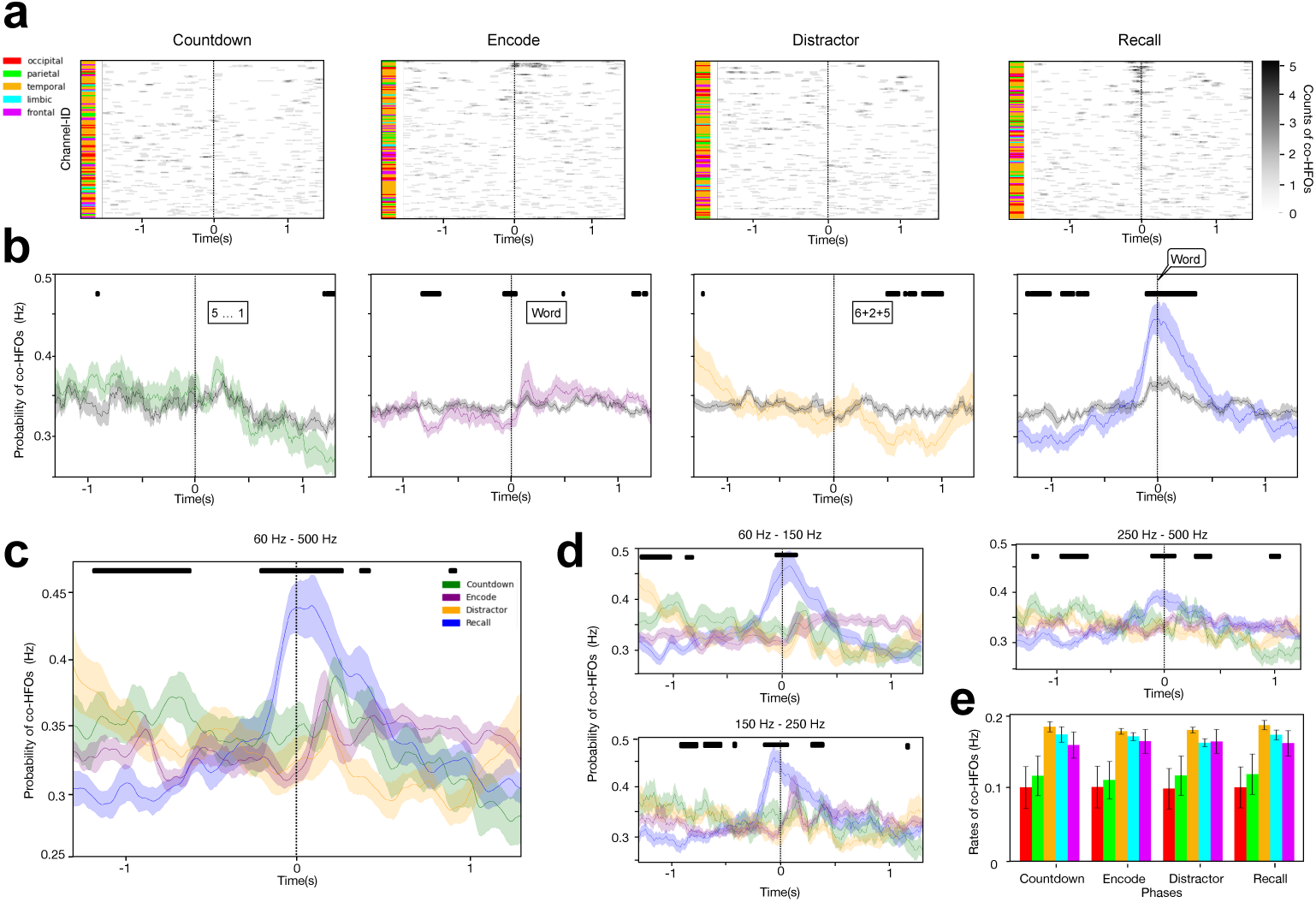
Coincident bursting across sensory and associational cortical areas is modulated by memory processing. a) Matrices of coincident HFO detections from an example patient, plotted as in Fig. 1c, show all channels localized in the five cortical lobes studied in a descending order from the most correlated pairs on top (Pearson’s correlation > 0.5). b) Probabilities of co-HFO bursting across the correlated channels from all patients reveal times of significantly increased or decreased co-activity relative to the other non-correlated channels (asterisks denote a significant difference). c) Comparison between the four task phases indicates significantly decreased probability of coincident HFO bursting approx. one second before recall verbalization (blue) with subsequent gradual rise and significantly increased probability a couple of hundred milliseconds before and after the verbalization onset (asterisks denote a significant difference between the four task phases). d) A consistent temporal pattern of HFO co-bursting is present in the high gamma, ripple and fast ripple frequency ranges. e) Summary of cortical lobe engagement in HFO co-bursting shows equal contribution in all task phases. * indicates p<0.05.

In each phase of the FR task, we observed global HFO discharges comprising coincident detections across all five cortical lobes (Fig. 2a). These coincident detections were more frequently observed around stimulus presentation and especially at the onset of free recall (Fig. 2b) on electrode pairs showing significantly correlated HFO bursting (Pearson’s co. relation, r > 0.50). The probability of detecting this co-HFO bursting differed significantly across the four task phases (One-way ANOVA: F(3,44) = 97.32, p = 3.4 × 10^-60^, N = 12). Specifically, this probability was significantly higher at the time bin of recall onset compared to the time bin of countdown, distractor and encoding onsets (One-way ANOVA, F(3,44) = 63.29, p = 6.88 × 10^-40^, N=12). Conversely, the probability was significantly lower in the time bin one second before recall (One-way ANOVA, F(3,44) = 2.21, p =0.013, N=12) and increased sharply ∼300 ms prior to reaching its peak at onset of vocalization Fig. 2c). During the encoding phase, the relative rate of co-HFO bursting was the highest during time of word presentation, likely reflecting active memory processing of the word displayed for 1.5 s on the screen. Although countdown digits and arithmetic equations were also presented for at least one second, they elicited significantly lower co-HFO bursting rates than word presentation during the one second period following stimulus presentation (One-way ANOVA, F(2,33) = 12.28, p = 5.56 × 10⁻^6^, N=12). These temporal patterns were consistent across the entire frequency range studied (60–500 Hz) as confirmed in three independent subranges (Fig. 2c).

No significant differences in the average rates of co-HFO bursting in particular cortical lobes were observed across the four task phases (One-way ANOVA: temporal: F(3, 40) = 0.45, p = 0.72; frontal: F(3, 40) = 0.02, p = 0.997; parietal: F(3, 40) = 0.02, p = 0.997; occipital: F(3, 40) = 0.001, p = 0.999; limbic: F(3, 40) = 0.45, p = 0.72; Fig. 2e). However, there was a significant effect of the cortical lobes on co-HFO bursting (one-way ANOVA: countdown: F(4, 50) = 2.90, p = 0.031; encode: F(4, 50) = 3.29, p = 0.018; distractor: F(4, 50) = 2.95, p = 0.029; recall: F(4, 50) = 3.10, p = 0.024), revealing a consistent pattern in all four task phases of lower rates in the occipital and parietal areas compared to the other three lobes studied (Fig. 2e). Altogether, these findings suggest that while co-HFO bursting occurs throughout all task phases, it is differentially modulated by memory processing with a relatively elevated rate during stimulus encoding, and a sharp peak in the probability of co-HFO bursting at the time of memory recall.

We tested further whether this modulation of coincident HFO bursting is related to memory or other sensory or motor processes. For instance, the peak of co-HFO bursting around the onset of verbalizing the recalled word could potentially be related to preparatory motor activity underlying vocalization. Notably we observed that the increased probability of coincident bursting preceded the very onset of vocalization by approximately 300 ms (Fig. 2c), resembling the timing reported of the readiness potential before subjective urge or will to make a movement ^38^. In one of the additional tasks, we used the same word pool in a cued recall PAL paradigm, where the patients were encoding 6 pairs of words instead of 12 words presented sequentially on the screen (Fig. 3a). The same subjects also performed an audio version of both the FR and the PAL tasks to ensure that the observed patterns were not specific to visual stimulation. During the PAL recall phase, only one word from each pair was presented as a cue for recalling its associate. In this paradigm we observed that the peak of co-HFO bursting shifted earlier, occurring around cue presentation approximately one second before the onset of vocalization (Fig. 3c). These finding demonstrated that increased co-HFO bursting (1) was related to memory retrieval triggered by the cue, (2) was not related to preparatory motor activity for word vocalization, and (3) was not specific to free recall, although the peak was more prominent in the FR task.

**Figure 3.**
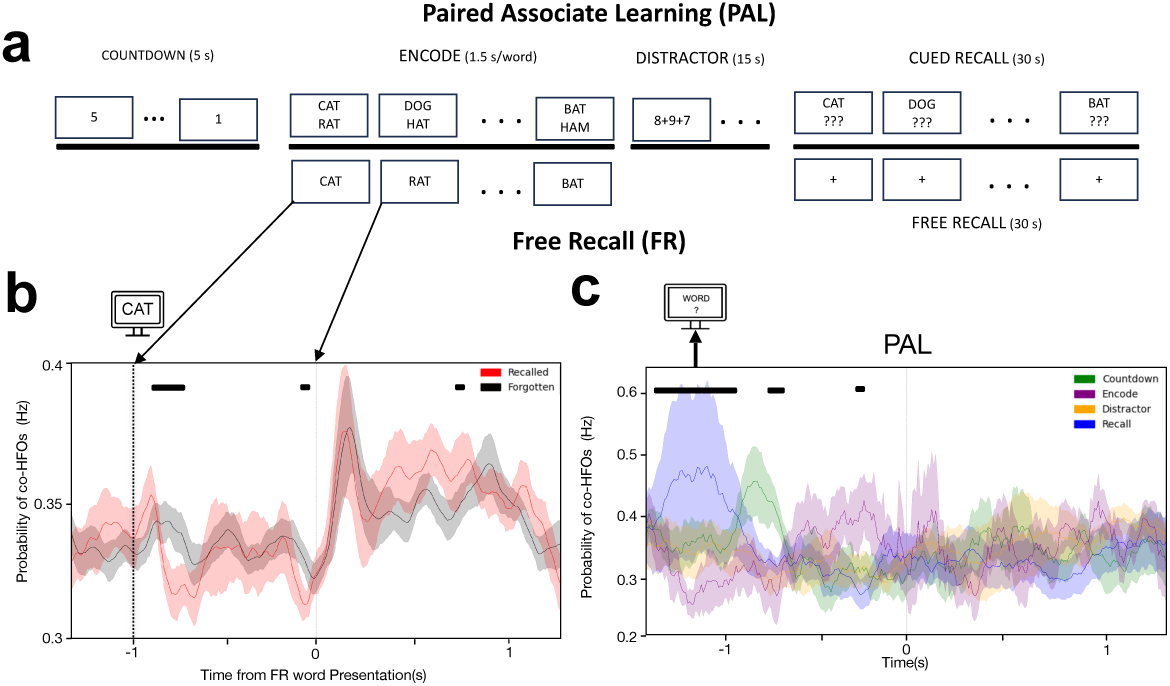
Memory processing modulates coincident HFO bursting during stimulus encoding and cued recall. a) Schematic of a cued-recall version of the task (top), alongside schematics of the free recall (bottom) all using the same pool of words. b) Probability of co-HFO bursting across the correlated channels from all patients is lower before and higher during presentation of subsequently recalled (red) compared to forgotten (black) words (black bar indicates significant bins at p<0.05). Notice the significant difference right before onset and following offset of the presented words on the screen, suggesting that suppression of co-HFO bursting during inter-stimulus interval is predictive of successful encoding. c) Analogous probability plots from the cued recall version of the task reveal enhanced co- HFO bursting at the time of the cue presentation and not at the onset of recall vocalization itself. Notice that the peak of co-HFO bursting occurs much earlier than in the free recall version of the task when the pair associated words are recalled, separated from any vocalization (black bar indicates significant difference between the task phases at p<0.05, as in Fig. 2c).

We also compared the co-HFO bursting probabilities during encoding between the words that were subsequently recalled and forgotten. The probability was lower during the inter-stimulus interval, especially after presentation of the previous word and before the moment of presenting words that were later successfully recalled (Fig. 3b). Conversely, It was relatively higher during presentation of the subsequently recalled words on the screen (Fig. 3b). These subsequent memory effects were observed not only in the grand-average analysis involving all cortical lobes, but also within individual lobes studied in both hemispheres (Suppl. fig. S4). This pattern suggests that suppression of coincident bursting in preparation for encoding new stimuli, followed by enhanced bursting during word presentation, heralded successful subsequent recall, confirming a role for co-HFOs in memory processing.

Given that the global co-HFO bursting across the cortex is modulated by memory processing, we investigated the proportion of areas engaged in encoding and recalling individual words (Fig. 4a). On average, 45.2% of all implanted contacts were involved in coincident HFO bursting during word recall, and 42.0% during word encoding. There were no significant differences in these proportions between the encoding and recalling phases (Chi-squared test, χ²(1)= 0.70, p = 0.999, N=12). Similarly, all five cortical lobes studied participated in co-HFO bursting in similar proportions (Fig. 4b) with no significant differences (Chi-squared test, χ²(4) = 0.56, p = 0.97, N=12). Thus, both sensory and associational cortical areas were equally involved in the co-HFO discharges.

**Figure 4.**
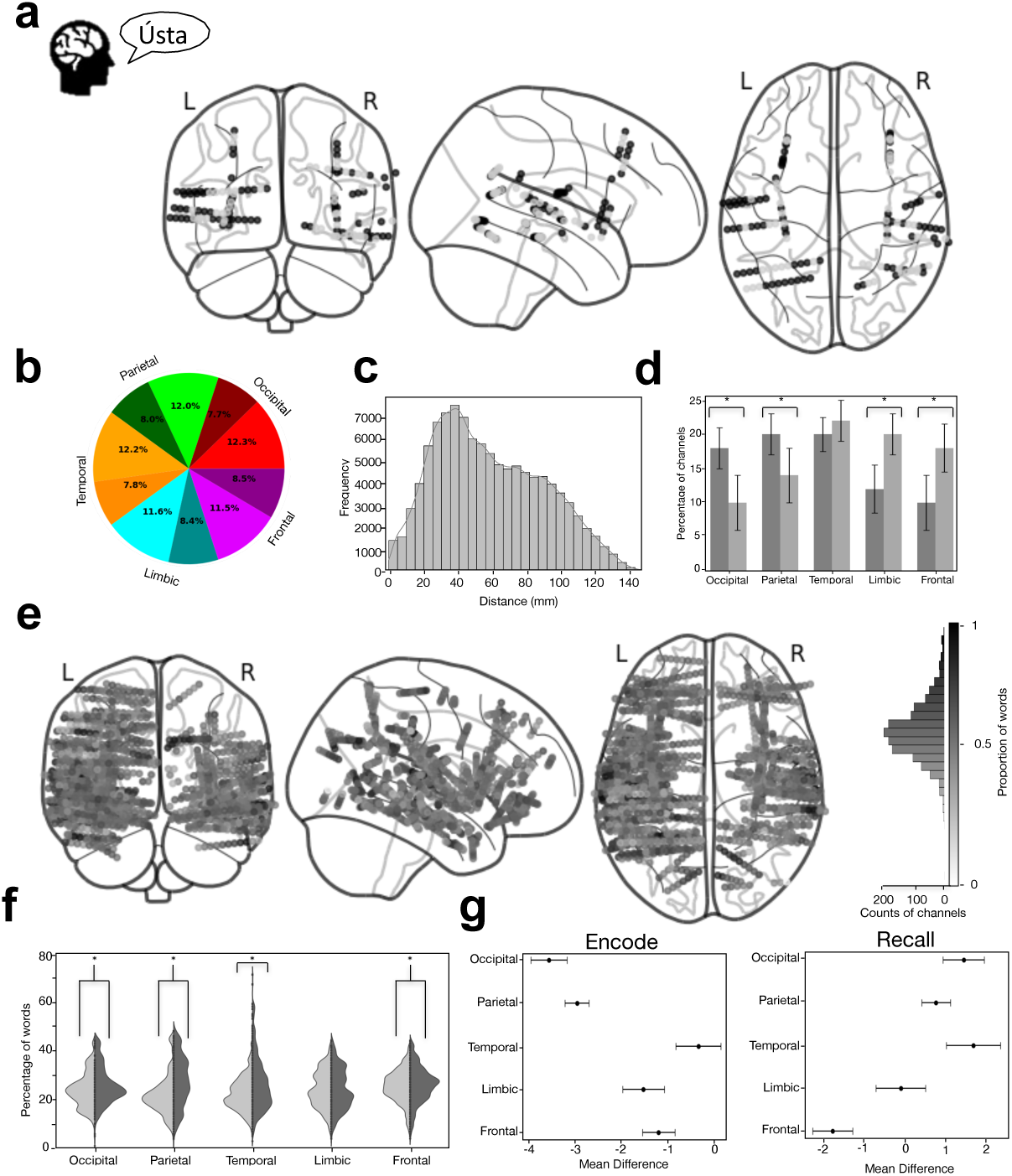
Widely distributed networks of coincident bursting are involved in encoding and recalling words. a) Localization of electrode channels that displayed coincident HFO bursting (black dots) during recall of an example word is distributed throughout the cortex engaging approx. half of the contacts implanted. b) Proportion of all channels showing significantly correlated (bright shades) and non-correlated (dark shades) co-HFO bursting averaged across all patients reveals equal contribution of all five lobes (color-coded as in Fig. 2). c) Anatomical distances between the correlated electrode channels (Pearson’s correlation, r > 0.50) show greatest density between 20-100 mm. d) Distribution of the correlated channels in the five cortical lobes is significantly higher during the encoding phase in more sensory areas of the occipital and parietal lobes, and higher during the recall phase in more associational areas of the limbic and frontal lobes. e) Summary of percent involvement of any one electrode contact (dot) in co-HFO bursting across the entire pool of words reveals relatively low specificity with most channels engaged in more than of the words recalled. Notice that most channels are involved in recalling a subset of words in the pool with only a few involved in almost every word (darker shades of gray). f) Violin plots summarize the average proportions of encoded (light gray) and recalled (dark gray) words that elicited co-HFO bursting of any one channel in a given lobe. g) Tukey’s post-hoc analysis compares the mean proportions from ‘f’ across the five cortical lobes in the two task phases. Notice that despite significant differences between the phases and the cortical lobes, the overall proportions are consistent ranging between approx. 10-50% of all words. * indicates p < 0.05.

This significantly coordinated bursting was not restricted to local channels within a single cortical lobe; most of the significantly correlated channels were separated by 20–100 mm (Fig. 4c), confirming widespread coordination over distances as large as 200 mm ^11^. Interestingly, despite the widespread engagement across cortical lobes in co-HFO bursting, we observed a significantly greater proportion of channels in the occipital (Wilcoxon signed-rank test, W = 10, N=5, p = 0.045) and parietal (Wilcoxon signed-rank test, W = 10, N=7, p = 0.035) lobes during the encoding phase compared to the recall phase. Conversely, the frontal (paired t-test, t(9) = 2.89, N=10, p = 0.014) and limbic (Wilcoxon signed-rank test, W = 14, N=12, p = 0.040) lobes exhibited significantly greater channel involvement during the recall phase (Fig. 4D). This differential activation reflects the expected contributions of more sensory and more associational cortical areas in the two task phases, with early processing regions more engaged in memory encoding and higher-order cognitive areas more engaged in memory retrieval.

How specific are these globally coordinated cortical networks to particular stimuli? In other words, are the same networks engaged in all instances of memory encoding and recalling or do they reveal specificity to particular words? We found that the majority of electrode contacts implanted participated in co-HFO bursting during the encoding or recall of approximately half of the words studied (Fig. 4e). These electrode contacts participated in varying proportions to the encoding and recall of words across the five cortical lobes, with significant differences in four out of the five lobes. The parietal lobe showed the largest difference in the proportion of words between the encoding and recall phases (t-test, t = -26.84, N=12, p = 2.05 × 10^-151, Cohen’s d = 0.63), followed by the temporal (t = -40.50, N=12, p < 0.001, d = 0.47), occipital (t = -10.09, N=12 p = 1.06 × 10^-23, d = -0.29), and frontal (t = -7.07, N=12 p = 1.56 × 10^-12, d = -0.11) lobes (Fig. 4f). There was a considerable overlap between the contacts engaged during encoding and recall of the same words, involving on average more than 10% of all contacts (Suppl. Fig. 5). Despite significant differences between the cortical lobes in both phases, the overall proportions varied within a consistent range of approximately 10-50% of words in the pool for any one contact (Fig. 4g). Hence, at least on the level of macro-contacts used in this study, the global co-HFO bursting patterns were not specific to individual word stimuli but rather showed a general engagement during encoding and recall of the words.

Finally, we asked whether the global co-HFO bursting during encoding or recalling a word is a unitary event or involves multiple related bursting events. Hierarchical clustering of the trains of HFO bursting from all electrode channels according to how similar they are across time revealed a temporal sequence of multiple bursting events, which was not visible when the contact channels were sorted according to the pairwise correlations. The clustering revealed multiple distinct networks of contacts bursting in a sequence - one cluster following another in time from the onset to the end of word recall (Fig. 5a). Each cluster comprised contacts from multiple cortical lobes and was related to other clusters in a complex tree hierarchy of correlation. Particular clusters showed enhanced co-HFO bursting at distinct moments in time each preceded by a suppression around 1s earlier (Fig. 5a), as shown in Fig. 2c. This temporal pattern of relatively suppressed co-HFO bursting before recall and induced bursting at the time of recall followed by a long ‘tail’ of elevated bursting probability fits the profile that we originally observed in the recall phase (Fig. 5a).

**Figure 5.**
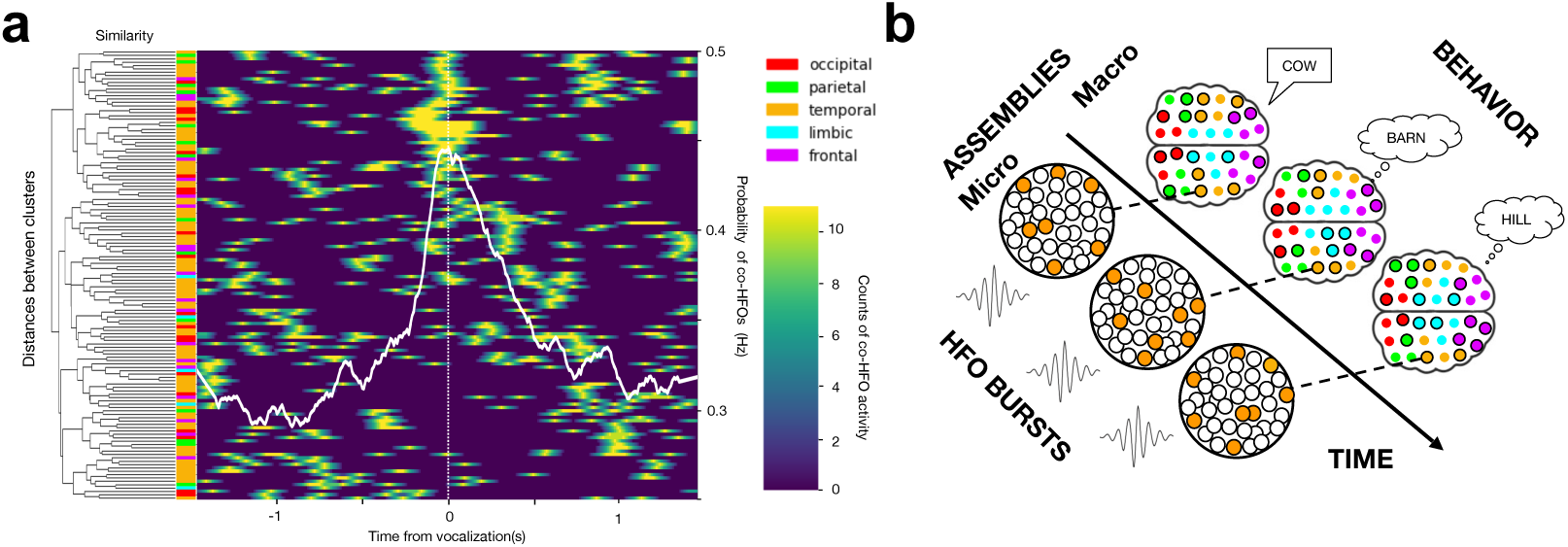
Global cortical bursting is temporally organized into sequences underlying word recall. a) Heatmap of hierarchical clustering of co-HFO bursting during recall of an example word reveals distinct sequential discharges, each involving multiple cortical lobes and aligning with the overlaid grand-average pattern of co-HFO probability from Fig. 3c (white line). Notice that the sequence of discharges along the diagonal of the heatmap is preceded by a parallel diagonal of co-HFO bursting suppression approx. 1 s before each discharge. b) Schematic of the model illustrates how local neuronal assemblies that generate the global sequences of co-HFO bursting can account for recall of a word followed by related concepts.

This successive propagation of co-bursting cortical networks is reminiscent of sequences in place cell firing reported in the rodent studies ^39–42^. In the context of our task, one bursting event underlying recall of a word could elicit another event related to recalling associated concepts (Fig. 5b). This would give rise to a sequence of successive co-HFO bursting events. We have recently proposed such sequential discharges of cortical HFO bursting on macro- and micro-scales for tracking electrophysiological engram activities in humans ^20^.

## Discussion

In this study, we show that bursts of high frequency oscillations co-occur over a large anatomical scale of multiple sensory and associational cortical areas. These coincident bursting happens across all phases of performing a task, but their occurrence is strongly modulated by memory recalling and encoding. This co-HFO bursting is suppressed before and enhanced during presentation of the stimuli that will subsequently be recalled compared to those that will be forgotten, and peaks approximately 300 ms before the recall vocalization. Around half of the recorded sites from all cortical lobes are engaged in this global coordination of HFO bursting underlying encoding and recalling the remembered stimuli, which was not specific to particular areas or words. Multiple distinct cortical networks were coordinated together in a sequence of bursting when recalling a single word. Instead of one bursting network, our results suggest a sequence of temporally coordinated networks. These findings suggest possible roles of globally coordinated discharges of high frequency oscillations in organizing communication and integration of information for memory processing.

Globally coordinated HFO bursting can be particularly important for memory processing, even though it is ubiquitous throughout all memory and non-memory related task phases. Previous studies found that ripple HFOs co-occur during sleep and waking periods but are especially prominent in memory recall ^11,17^. Our results confirm the maximum occurrence of coincident HFO bursting around the onset of recall vocalization. Interestingly, the significantly greater occurrence compared to stimuli presentations in the other task phases started 300 ms before the vocalization onset and was significantly suppressed approximately 1 s before the onset (Fig. 2c). This temporal timeframe is in agreement with the expected dynamics of recalling an item - first a suppression of bursting in response to the previously recalled item would be followed by a gradual build-up of activity corresponding to recalling a new item. The timing resembles the ‘readiness potential’ from Libet’s original experiments ^38^, which was more recently also demonstrated at the level of single unit firing ^43^. While this study was not designed to address the emergence of conscious volition or its neural substrates ^44^, the timing of this peak in coincident HFO bursting preceding free recall at least suggests that it is related to memory processing and not to other processes like preparation or planning for vocalization. This was confirmed in the additional cued recall paradigm, which showed a shift of the peak co-HFO bursting to 1000 ms before word vocalization (Fig. 3a) around the time of cue presentation.

Furthermore, it was not only memory recall but also memory encoding that induced more coincident HFO bursting after stimulus presentation than in the other task phases. There was an increased occurrence starting with a local peak at 150-200 ms after stimulus onset followed by an elevated rate throughout the time of word presentation on the screen, which corresponds to the expected information processing timeframe that we observed in the same paradigm across the ventral visual stream ^45,46^. Like in the recall phase, the increased occurrence was preceded by a suppression in co-HFO bursting with one dip following presentation of the previous word and another right before onset of a new word presentation. Word stimuli that showed less of this suppression before and less enhancement after presentation were more likely to be subsequently forgotten (see Fig. 3b). Altogether, these results point to an important role of the globally coordinated HFO discharges not only around memory recall but more generally in memory processing.

Previous studies proposed an even more general role for the global HFO bursting in integrating information processing across the brain ^17,30^, which would provide a solution for the binding problem ^1,8,47^. Such a role would apply both to memory, attention and perceptual functions ^2,5,7,48^. In agreement with a more general role, our results show internally induced co-HFO bursting in the occipital cortex during recall in the absence of any external visual stimulation (see Fig. 1). On the other hand, visual stimulation with countdown digit or algebra equation presentations during the preparatory and the delay phases of the task, respectively, induced a smaller increase in co-HFO occurrence (see Fig. 2b), even though both were perceived and engaged subjects’ attention in the task. One could argue that these other less relevant stimuli were not as attended or requiring as much cognitive effort as memorizing or recalling items. In other words, perceiving countdown digits or solving algebra is more automated compared to intentionally focused formation of new memory traces and especially their internally-driven retrieval, which would arguably require more of global cortical coordination. It would explain why cued recall induced a smaller rise in the co-HFO bursting than the free recall (see Fig. 3a). Dissociating specific roles in particular cognitive functions would necessitate a more appropriate experimental paradigm.

Still, high frequency oscillations especially in the ripple frequency range have been specifically associated with memory processing given the link with hippocampal sharp-wave ripple complexes ^10,12,13,18,28,29,32,49,50^. Bursts of oscillations in gamma and ripple frequencies are ideally suited for inducing synaptic plasticity ^51^ and transferring information in ‘packets’ across the cortex ^52^, congruent with the proposed roles in integrating and storing information in memory. In human studies, identifying and defining a particular electrophysiological activity corresponding to the rodent sharp-wave ripple complexes is challenging due to lack of clear detection criteria ^20,24^, including precise anatomical localization, presence of a sharp wave in addition to an oscillation above a particular power threshold and a minimum number of cycles in the unfiltered signal. Hence, based on our previous studies ^13,14^ we took a more general approach detecting distinct oscillations across a broad frequency range and all cortical locations with no need for isolating hippocampal sharp-wave ripple complexes. Rodent studies showed that high gamma/epsilon and ripple oscillations in the hippocampus shared common neuronal ensembles and mechanisms ^21,53,54^, despite clear differences even among ripple detections ^55^. Using a similar threshold for detecting oscillations of at least four cycles, we found that the global co-HFO bursting is not limited to a narrow ‘ripple’ frequency range but involves oscillations across gamma, ripple and fast ripple ranges. The same pattern of induced coincident bursting before recall onset was present in 70-150Hz, 150-250 Hz and even 250-500 Hz, although the peak was gradually fading with increasing frequency of the detected bursts (see Fig. 2c). Interestingly, at these higher frequencies there is an increased likelihood of detecting pathological HFOs in this patient population ^56^ (our analyses were limited to HFO detections, which are temporally related to memory processes in the tasks). The reported patterns of co-HFO bursting were also not any stronger in the limbic cortex, which included the hippocampus, than in the other cortical lobes (see Suppl. fig. 4). Therefore, it is plausible that the global cortical bursting may have a more general role than the one previously proposed for hippocampal-cortical dialogue in human memory encoding, consolidation and retrieval ^12,15,16,18^.

We have limited our study to bursts of high-frequency oscillations above the classic Berger bands. Similar bursts, however, can be detected in the lower frequencies. A recent study by Sieber et al. reported hippocampal ‘bouts’ of theta frequency oscillations preceding a turn in real-world or imaginary navigation in epilepsy patients ^57^. The theta ‘bouts’ were on average four cycles long, they were not related to motor activities when making a turn, and were used to reconstruct spatial locations even in the imaginary navigation. Similar bouts, discharges or bursts of approximately four cycles of oscillations were also found in alpha and beta frequencies in that study, which suggest a universal organization of cortical oscillations. How the high and the low frequency activities are related to each other in time and anatomical space ^46,58,59^ remains to be explored on the level of individual oscillatory discharges, but a common role emerges. Whether it is freely recalling word concepts or imagining spatial locations, these abstract cognitive processes are preceded by the discharges of low and high frequency oscillations.

What do these global coincident bursts reflect? We have recently proposed HFOs as electrophysiological biomarkers of engram activities ^20^. Individual bursts are generated by neuronal assemblies that coordinate their firing when processing information about a particular item or its features. Coincident bursting across the cortex can integrate this processing in large scale networks of assemblies processing features of related items. Our results suggest that these large-scale networks of assemblies underlying encoding or recall of any one item are widespread, involve around half of the recording channels, and are not highly specific to any particular words (see Fig. 4). This picture is similar to a map of all cells activated in response to encoding or recalling a single memory trace - an engram of episodic memory in mice, which engaged up to half of all the brain regions studied ^35^. To spot differences between such large-scale maps of two separate engrams for episodic memory traces would be possible on the level of single activated cells but challenging on the level of entire brain regions. That is likely why our macro-contact recordings resulted in responses to a large proportion of word items and relatively high overlap between them. Using micro- or meso-contact recordings would be expected to resolve more specific engram contributions. There are probably multiple distinct neuronal assemblies generating HFOs corresponding to different words that would be detected on the same macro-contact or distinct meso- or micro-contacts. The recent study by Vaz and colleagues confirmed that detecting single neuron firing on micro-contacts in parallel to HFO bursts on macro-contacts can be used to decode correct and incorrect recall of specific words from sequences of single cell firing ^9^. Future studies employing combined macro- and micro-scale electrophysiological recordings ^23,60^ promise to track the hypothetical engram activities not only with high spatial resolution but also in time to follow dynamically changing neuronal assemblies ^61^. Direct electrical stimulation to evoke engram activities ^62,63^ at macro- and micro-scales offers causal evidence for these hypotheses.

Although our study was limited to macro-contact recordings, we also observed two levels of organization for coincident HFO bursting. On the first level, HFOs generated by local neuronal assemblies were coordinated across the cortex in global coincident bursting, as is now increasingly reported in the most recent studies ^15–17,19^. This level can basically be described as a global network of local assemblies. The second level that emerged from our results comprises a larger network of multiple distinct global bursting networks, which are coordinated together in a temporal sequence (see Fig. 5). This macro-scale ‘sequence of networks’ could be compared to the micro-scale ‘synfiring chains’ ^64^, in which firing trains of one neuron or one assembly of neurons corresponding to the first item triggers another train corresponding to the second item, then the third, and so on. In summary, there is a ‘micro’ level of coordinated sequential firing of neuronal assemblies giving rise to HFO bursts, and a ‘macro’ level of sequentially coincident bursting in a higher order global network organization, as we have recently proposed ^20^. The ‘macro’ level sequences of the global coincident bursting remain to be further investigated.

There are other outstanding questions about the global HFO bursting that need to be addressed in future studies. First of all, can this widespread temporal coordination of local neural activities provide a mechanism to address the binding problem ^1,47^? Recent studies suggest so ^11,17,30^ but testing it would require an appropriate behavioral paradigm probing conscious awareness of the perceived stimuli. Oscillations in the high frequency ranges (above 50 Hz) were shown to reflect explicit awareness of visual stimuli in monkeys ^65^ or conscious dream experiences in humans ^66^. In terms of these studies, coordination of HFOs across the brain in coincident bursts would provide a plausible neural mechanism for concepts like ‘ignition’ or ‘global broadcasting’ in the global neuronal workspace hypothesis ^67^. One could then ask whether the sequences of bursting events correspond to a train of thought or a stream of consciousness. In the case of our paradigm, we can only speculate about such epiphenomena; the sequences reported can, however, be explained more concretely in terms of a model for recalling engram representations of related word concepts from the memorized lists (see Fig. 5). Coincident HFO bursting presents a testable activity for these and other models of cognitive functions, including perceptual binding, conscious awareness or engram representations of related concepts in the human mind.

Having a concrete electrophysiological activity reflecting abstract processes in the human mind would enable not only tracking across the brain and time but also modulating them with targeted brain stimulation in brain disorders. Memory and cognitive deficits are a hallmark of neurological and psychiatric diseases with limited treatment options. Emerging technologies for chronic brain recording, analytics and stimulation using distributed cloud computation and machine learning tools can now deliver personalized adaptive modulation therapies for restoring movement ^68^, mood ^69^ and memory functions ^70^. Biomarkers of electrophysiological activities like the HFO bursting promise to guide, monitor and assess new neuromodulation therapies across the anatomical brain regions and time.

## Methods

### Participants and Electrode Localization

Seventeen patients (10 males, mean age 31.94 ± 1.63 years) undergoing intracranial stereo EEG seizure monitoring for epilepsy surgery at St. Anne’s University Hospital in Brno and Jan Mikulicz-Radecki University Hospital in Wroclaw were recruited in this study, which was approved by the local Institutional Review Board ethical committees(Table 1). Participants were native speakers of Czech (n=8), Slovak (n=4), or Polish (n=5), and all provided written informed consent (Table 1). Electrode implantation was dictated by clinical priority of treating drug-resistant seizures, leading to a non-uniform distribution of electrodes across all cortical areas (Suppl. table 1). Data from five patients were excluded from further analysis due to technical issues related to accurate synchronization of the task events with the electrophysiological recordings, which was critical for this study. Following implantation, pre-operative high-resolution CT and post-operative MRI scans were merged and normalized to determine electrode locations in MNI space and assign anatomical labels according to the Automated Anatomical Labelling atlas (AAL3) ^71^ . Electrodes were grouped into distinct cortical lobes: occipital, parietal, temporal, frontal and mesial temporal (referred to as ‘limbic’). The limbic lobe included hippocampus, amygdala, posterior cingulate cortex and parahippocampal cortex. In total, 3,239 recording contacts were implanted across all patients (2,957 macro contacts).

### Experimental design

Participants completed a battery of verbal memory tasks designed to examine neural dynamics underlying memory encoding, maintenance, and retrieval of words. The main Free Recall (FR) task involved presenting 12-word lists derived from common nouns in the participant’s native language (Czech, Slovak, or Polish), sourced from a publicly available repository (http://memory.psych.upenn.edu/WordPools). In total, 180 words from the pool were selected based on their translatability to the three Slavic languages, and randomly allocated to 15 lists before the start of a task run. Each run started with a 5 s countdown presentation of digits from 5 to 1 immediately followed by presentation of a word list for encoding. Each word from a list was displayed for 1500 ms with 1000 ms inter-stimulus interval. Following the final word, participants engaged in an arithmetic distractor task (e.g., A + B + C = ?, where A, B, and C were random integers between 1 and 9) to prevent rehearsing the list. After 20 s of the distractor, participants had 30 s to verbally recall as many words as possible in any order. Verbal responses were recorded with high-fidelity microphones and annotated for accuracy using an automated transcription method with supervision and post-hoc verification (Suppl. fig. 2). The transcription process utilized a custom interface with integrated fine-tuned Whisper model. To achieve precise word alignment, a timestamped modification was implemented using Dynamic Time Warping. This algorithm adjusted timestamps by matching predicted text with actual speech patterns, correcting timing discrepancies and improving synchronization.

Precise times of initiating the verbal responses were automatically marked by the transcription method and manually corrected if necessary (Suppl. fig. 2) to mark the very onset of the first sounds of word verbalization. All other task events were automatically time-stamped in the electrophysiological neural recordings via TTL pulse generation from the task computer.

In addition to the FR task, participants completed an analogous cued recall task called Paired-Associate Learning (PAL). Instead of the 12-word lists, 6 word pairs were sequentially presented one pair after another. After the same arithmetic distractor task as in FR, participants were presented with a cue of one word from each pair, prompting them to recall its corresponding paired associate. The PAL task also comprised 15 lists of randomized word pairs, which were taken from the same word pool as in the FR task. An additional audio version of the FR and PAL task was completed by some patients with the presented words and the recall cue word played out loud with a fixation cross on the computer screen.

### Electrophysiological Recordings

Electrophysiological recordings were performed using ATLAS neurophysiological system (FHC-Neuralynx Inc.) from up to 128 channels recorded simultaneously at 32 000 Hz sampling rate (Jan Mikulicz-Radecki University Hospital in Wroclaw), and BrainScope BioSDA09 system (M&I Ltd.) from up to 192 channels at 5 000 Hz sampling rate (St. Anne’s University Hospital in Brno). The signals were recorded from multiple standard depth electrodes (2.3 mm exposed surface, 5–10 mm inter-contact spacing) for stereo EEG surgical procedure (AD-Tech Inc.) implanted for prolonged seizure monitoring. Additional subgaleal contacts were used as reference electrodes. To enhance spatial specificity and minimize volume conduction artifacts, signals were re-referenced using a bipolar montage applied to adjacent contacts within each stereotactically implanted depth electrode.

All electrophysiological time-series data were stored in Multiscale Electrophysiology Format (MEF) version 3.6 and structured in accordance with the Brain Imaging Data Structure (BIDS) format, ensuring universal accessibility and reproducibility^36^. Participant data were anonymized with personally identifiable information removed in adherence to ethical guidelines. The complete dataset has been published and is available on the EBRAINS portal ^72^.

### Detection of individual HFO bursts

Individual bursts of the high-frequency oscillations (HFOs) were detected according to our previously developed methodology ^13,14^.The signals were first decomposed into 38 logarithmically spaced frequency bands ranging from 60 to 800 Hz using zero-phase finite impulse response filters with 200 ms transition bands and a −6 dB cutoff. The amplitude envelope for each band was extracted using the Hilbert transform and normalized via a sliding z-score transformation, with the mean and standard deviation computed over 10-second windows. Candidate HFOs were initially identified when the z-scored amplitude exceeded a liberal threshold of 2 standard deviations (SD) for at least three consecutive oscillations. Events spanning multiple frequency bands were merged, and for each detected event, the dominant frequency—defined as the band with the highest z-score—was determined. The maximum amplitude within the event window was also extracted to characterize peak activity.

To enhance specificity of the detections and remove artifacts, a dual-threshold detection approach was implemented. Only events with a peak z-score exceeding 3 SD were retained, and an additional duration criterion required that events persist for at least four complete oscillatory cycles at their dominant frequency. To further ensure that detected events represented true oscillatory phenomena rather than transient spectral fluctuations, a final cycle-count verification was performed, requiring at least one complete oscillation above the threshold for an event to be classified as an HFO burst. This approach was designed to reduce false positives in the human intracranial recordings, where greater artifact susceptibility necessitates more stringent detection criteria than those typically applied in rodent studies.

### Analysis of coincident HFO bursting

HFOs were represented as discrete points at the peak amplitude in frequency-time space ^37^, which implies the quantization of each burst into discrete events with intrinsic features such as onset time, offset time, maximum frequency, minimum frequency. The occurrence rate was computed for each phase (countdown, encode, distractor, recall) across three frequency ranges: 60–150 Hz, 150–250 Hz, and 250–500 Hz, (Suppl fig. 1). Raster matrices were constructed to visualize the temporal distribution of the HFO detection events, using a 10 ms bin size, selected based on a typical HFO duration of 10–50 ms. Time zero for each task phase was defined at digit presentation (countdown), word presentation (encode), arithmetic task onset (distractor), and onset of remembered word vocalization (recall). Rasters were aligned to these event times within a ±1.5 s window.

To assess spatiotemporal relationships, HFOs were binarized (1 for HFO presence, 0 for absence), and a ±50 ms window (±5 bins) were examined across channels to identify coincident HFO bursts (co-HFOs). A co-HFO was defined as an HFO occurring within a ±50 ms window of another HFO on a different channel, ensuring that only temporally proximate events were considered functionally related. Pearson correlation coefficients were computed between channels for each raster, classifying channels as correlated (r > 0.5) or non correlated (r ≤ 0.5) and channels were sorted into two groups based on the correlation values (Fig. 2a). This threshold was chosen to prioritize strong functional relationships while mitigating the impact of noise and weak correlations. Given the large number of channels per subject (≥100), an r > 0.5 threshold was chosen to ensure that only strong co-firing patterns were considered significant.

To quantify functional connectivity dynamics, channels were separated into positively correlated (r > 0.5) and non-correlated (r ≤ 0.5) groups, and the rate of co-HFOs per time bin was computed separately for each group. This approach allowed for a direct comparison of co-activation patterns between correlated and non-correlated channels over time (Fig. 2b). The analysis was performed by averaging co-HFO rates across all detected events within each task phase. To account for inter-subject variability, we first computed the mean co-HFO rate for each subject at each time bin. To enable standardized comparisons across subjects, we then applied probability normalization, ensuring that the sum of all co-HFO rates across all subjects and time bins equaled 1. This transformation yielded a traditional probability distribution, where each value represented the probability of a co-HFO occurring at a given time bin relative to the entire dataset. For visualization purposes, the normalized values were scaled by a factor of 100 to express probability per second in figures, given the 10 ms bin size. Importantly, the normalization was applied after rescaling to a per-second probability, allowing for standardized comparisons across subjects and task phases. The resulting probability of co-HFO occurrence provided a standardized measure of how co-HFOs evolved over time relative to task events (Fig. 2c).

We extended our coincidence analysis to compare the temporal dynamics of co-HFOs between subsequently recalled and forgotten words (Fig. 3b) and across task phases in the paired-associate learning (PAL) task (Fig. 3c). These comparisons allowed us to assess whether functional network engagement varied as a function of recall success and task structure. Regional specificity was further examined by assessing co-HFO activity within individual cortical lobes to determine whether co-HFOs followed regionally distinct activation patterns (Suppl. Fig. 4).

For each patient, we identified electrode contacts involved in correlated bursting using co-HFO matrices as in Fig. 2A. These rasters provided a structured output from burst detection analyses, listing implicated contacts and their corresponding cortical lobes. Contacts from correlated HFO bursts were extracted and assigned lobe identities using the Automated Anatomical Labeling Atlas (AAL3) ^71^, mapping each contact to its respective cortical region. Thus, impact of pathological epileptiform HFOs was minimised by including only channels that correlated their bursting in response to a cognitive task event.

### Anatomical distribution and stimulus specificity analyses

To assess spatial distribution of the electrode contacts showing significantly correlated HFO bursting during memory processing, the brain coordinates of every contact were mapped and grouped into the five cortical lobes studied (Fig 4a). Euclidean distances between any two correlated channels were calculated and summarized as histograms to characterize the spatial extent of coincident HFO bursting (Fig. 4c).

For each lobe, we calculated the proportion of correlated channels as the ratio of correlated channels to the total implanted channels in that lobe, with non-correlated proportions derived by subtraction (1 − correlated proportion). These subject-level proportions were pooled across participants and averaged to obtain group-level estimates. To account for inter-subject variability in electrode coverage, we normalized the final proportions by expressing the number of correlated and non-correlated channels as a percentage of the total implanted channels in each lobe, ensuring that their relative distributions were preserved for group-level comparisons (Fig. 4b).

Coordinates were anatomically normalized using the automated anatomical labeling atlas (AAL3) ^71^ to account for inter-subject cortical variability. The proportion of words in recall in which a given electrode contact exhibited significant HFO bursting were pooled across subjects and mapped spatially (Fig. 4e). To quantify the engagement of each channel, we computed the frequency of significant co-HFO bursting occurrences across all trials in which a word was encoded or recalled. This provided a measure of how many words a given contact responded to with significant coincident bursting. This frequency for each contact was determined using a group-wise count transformation, which aggregates per-channel engagement occurrences across trials within each patient before pooling across the cohort. Specifically, for each electrode contact, we first counted the number of trials where significant co-HFO bursting was observed. These counts were then summed separately for encoding and recall within each participant. Finally, to generate group-level engagement distributions, individual patient results were pooled while normalizing for inter-subject variability by expressing each subject’s data as a proportion of their total implanted channels, thereby ensuring comparability across participants variability in total electrode contacts implanted.

To obtain cohort-wide estimates, per-patient engagement frequencies were concatenated into two aggregated datasets: one for encoding and one for recall. These datasets were then filtered into their respective lobes. Violin plots were generated for each lobe to compare engagement frequency distributions between encoding and recall. Each plot displayed the density of bursting occurrences per lobe, individual data points within each lobe group, and statistical significance markers for between-phase comparisons.

### Hierarchical Clustering Analysis

To identify patterns of co-occurrence across channels, we applied hierarchical clustering to the co-HFO matrices. The first step involved computing a correlation distance matrix, where the distance between two channels was defined as one minus the Pearson correlation coefficient. This metric ensures that channels with highly similar activity patterns (highly correlated signals) are considered closer, while those with dissimilar activity are treated as farther apart.

We then performed agglomerative hierarchical clustering using the average linkage method. In this approach, clusters are merged iteratively based on the average pairwise distance between all elements in the respective clusters. Average linkage (also known as UPGMA – Unweighted Pair Group Method with Arithmetic Mean) is particularly effective in preserving local structures while still capturing broader functional relationships between channels. Compared to other linkage methods, such as single linkage (which is sensitive to chaining effects) and complete linkage (which emphasizes compact clusters), average linkage provides a balance between local and global structure retention.

The hierarchical clustering results were visualized using a clustermap, where channels were reordered based on the clustering hierarchy. This reordering enhances the interpretability of the co-HFO matrices by grouping functionally similar channels together, revealing spatially and temporally coherent network patterns. All analyses were conducted in Python using NumPy, Pandas, SciPy, and Seaborn.

### Statistical Analysis

All statistical analyses were conducted using Python to ensure rigorous and reproducible evaluation of the data. The normality of the data distributions was first assessed using the Shapiro-Wilk test (α = 0.05). This test was employed to determine whether parametric (e.g., ANOVA, t-tests) or non-parametric (e.g., Kruskal-Wallis, Wilcoxon signed-rank) statistical approaches were appropriate for subsequent analyses.

To compare HFO bursting rates and coincident bursting probabilities across the four task phases (Countdown, Encoding, Distractor, Recall), we used the Kruskal-Wallis test, a non-parametric alternative to ANOVA that does not assume normally distributed data. Post-hoc comparisons between specific phases were conducted using Tukey’s Honest Significant Difference (HSD) test, with a family-wise error rate (FWER) of 0.05, ensuring control over multiple comparisons. For temporal dynamics of coincident bursting, one-way ANOVA was applied to detect significant differences in bursting probabilities over time. Tukey’s HSD post-hoc test was used to identify specific time points that contributed to significant effects. To examine the spatial distribution of co-HFO bursting across different cortical lobes, we applied Pearson’s Chi-squared test, separately analyzing encoding and recall phases. This test determined whether different cortical regions were equally involved in coordinated HFO bursting or whether specific lobes were preferentially engaged. To assess whether the same cortical regions were engaged during encoding and recall within individual subjects, we performed Wilcoxon signed-rank tests for non-normally distributed data or paired t-tests for normally distributed data. These tests helped to determine if a shift in co-HFO involvement occurred between encoding and recall. To investigate the subsequent memory effect, comparing HFO bursting rates between words that were later successfully recalled versus forgotten, we employed independent t-tests. This allowed us to evaluate whether memory-related differences in neural activity could be detected during initial encoding. To identify significantly co-activated electrode channels, we performed Pearson correlation analysis (r > 0.5). This method quantified the degree of synchronization between electrode contacts, revealing functional connectivity patterns during memory processing. For evaluating the spatial clustering of co-HFO bursting, we conducted permutation testing (10,000 iterations, FWER-corrected). This non-parametric approach tested whether observed clustering patterns exceeded those expected by chance, strengthening the reliability of spatial coordination findings .Finally, effect sizes for key comparisons were quantified using Cohen’s d, with 95% confidence intervals (CIs) to provide standardized measures of observed differences. All statistical tests were two-tailed and results were reported as mean ± standard error of the mean (SEM) unless stated otherwise.

