## Supplementary Material for "Coincident bursts of high frequency oscillations across the human cortex coordinate large-scale memory processing"

**Table S1. Distribution of electrode contacts implanted across cortical areas for treatment of drug-resistant epilepsy treatment.** Electrode placement varied across cortical regions based on clinical priorities, leading to non-uniform coverage across patients. The table summarizes the number of electrodes implanted per structure from all patients, reflecting variability in cortical sampling due to patient-specific clinical needs.

| Structure | N electrodes |
|---|---|
| Amygdala | 32 |
| Angular gyrus | 24 |
| Calcarine fissure | 4 |
| Caudate nucleus | 4 |
| Cingulate gyrus | 35 |
| Cuneus | 11 |
| Fusiform gyrus | 40 |
| Gyrus rectus | 17 |
| Heschl's gyrus | 10 |
| Hippocampus | 156 |
| Inferior frontal gyrus | 104 |
| Inferior occipital gyrus | 2 |
| Inferior parietal gyrus | 38 |
| Inferior temporal gyrus | 103 |
| Insula | 172 |
| Lingual gyrus | 31 |
| Middle occipital gyrus | 19 |
| Middle temporal gyrus | 313 |
| Orbitofrontal cortex | 56 |
| Paracentral lobule | 4 |
| Parahippocampal gyrus | 27 |
| Postcentral gyrus | 47 |
| Precentral gyrus | 38 |
| Precuneus | 37 |
| Putamen | 49 |
| Rolandic operculum | 42 |
| Superior frontal gyrus | 52 |
| Superior occipital gyrus | 10 |
| Superior parietal gyrus | 14 |
| Superior temporal gyrus | 124 |
| Supramarginal gyrus | 61 |
| Temporal Pole | 41 |

**Figure S1. Detection of high-frequency oscillations (HFOs) in intracranial recordings.** Raw signal (top) is presented with its decomposition into example ranges of gamma (60–80 Hz), ripple (80–250 Hz), and fast ripple (250–600 Hz) frequency bands. The amplitude envelope was extracted using the Hilbert transform and z-scored. The bottom panel displays the Hilbert matrix, illustrating spectral power dynamics over time. HFO bursts were identified based on amplitude thresholds and cycle-count criteria to distinguish true oscillatory events from transient spectral-power artifacts.

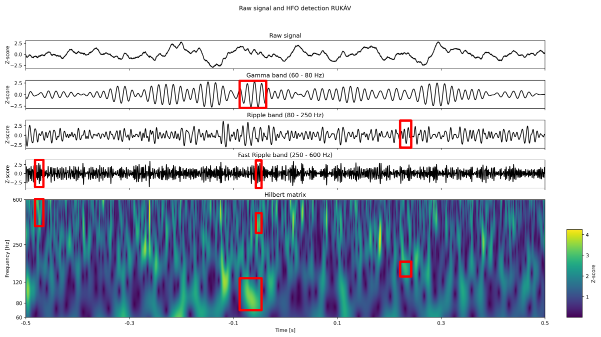

**Figure S2 Dynamic Time Warping (DTW) alignment of verbal responses with audio processing.** (a) Example of one DTW-based alignment of spoken words with their corresponding timestamps (red lines) is presented on a spectrogram of the recorded patient audio responses. The bottom panel presents the Mel-Frequency Cepstral Coefficients (MFCC) spectrogram of the audio signal. (b) Verbal responses were first automatically annotated using a fine-tuned Whisper model and then manually supervised and corrected, if needed, with timestamp realignment to ensure precise onset of the very beginning of word vocalization.

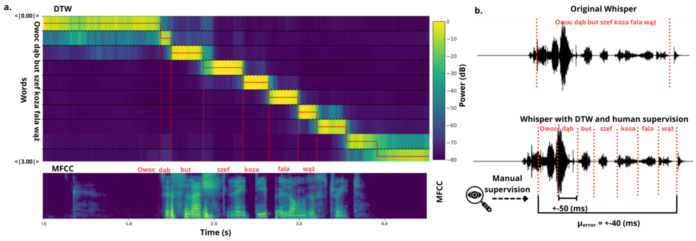

**Figure S3. Baseline HFO bursting rates show no differences across task phases.** Bursting rates per frequency range (60–150 Hz, 150–250 Hz, and 250–500 Hz) across the color-coded task phases reveal no significant differences (Kruskal-Wallis test, $H(3) = 2.94$, $p = 0.40$, $N = 12$) and consistent rates across patients (points) in all three frequency ranges. Error bars indicate standard deviation.

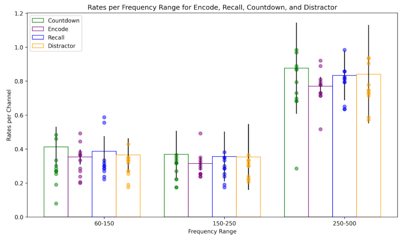

**Figure S4. Co-occurrence of HFO bursts across cortical regions shows consistent differences between trials with subsequently remembered and forgotten words.** Probabilities of co-HFO bursting, plotted as in Fig. 3b, for each lobe (row) and hemisphere (column) revealed the greatest differences (black arrows) in the right limbic and frontal lobes. Notice that in all cases the probability sharply decreases just before word presentation (time 0), followed by increased bursting during the time of word processing from around 150-1000 ms. Error area indicates the standard error of the mean.

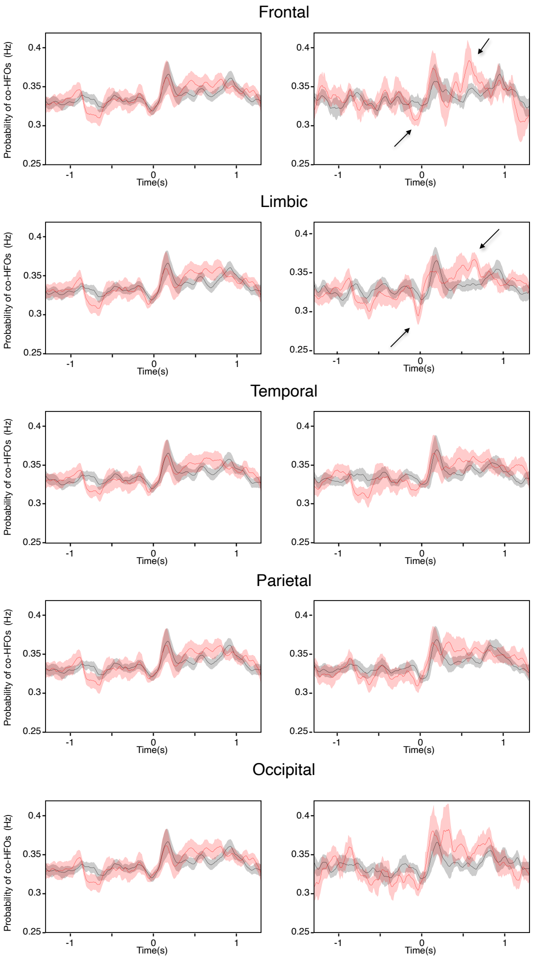

**Figure S5. Summary of the channel overlap during encoding and recall of the same word.** Bar plots show the proportion of channels exhibiting co-HFO bursting during encoding and recall of each word (x-axis) across the 12 subjects (each panel is one patient). Notice that despite wide variability in the proportion of overlapping channels and the words recalled, more than 10% of all contacts were on average engaged in co-HFO bursting during encoding and recall of the same word.

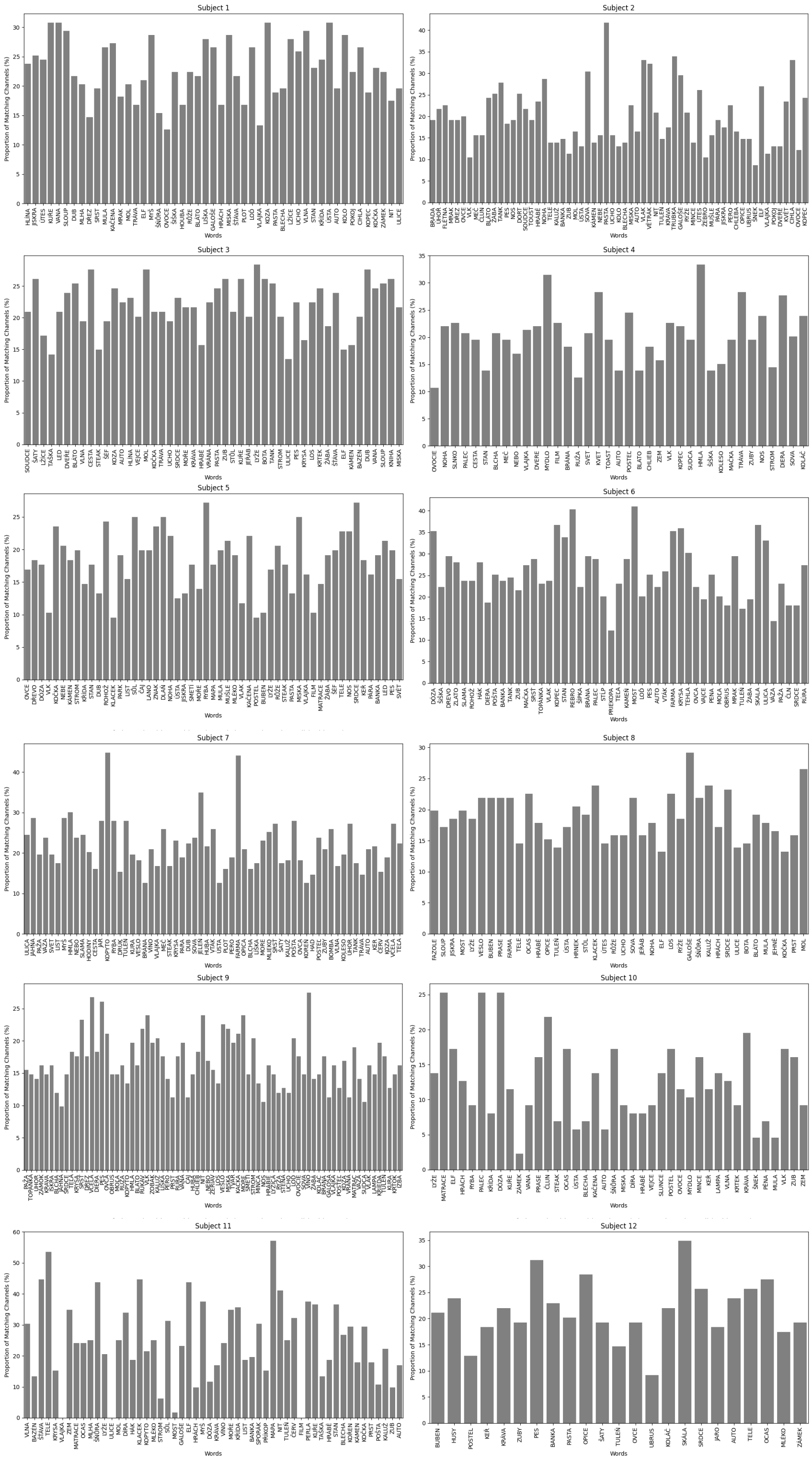
